# Regulatory crosstalk between motility and interbacterial communication in *Salmonella* Typhimurium

**DOI:** 10.1101/2020.09.09.289983

**Authors:** Jonathan Plitnick, Fabienne F.V. Chevance, Anne Stringer, Kelly T. Hughes, Joseph T. Wade

## Abstract

FliA is a broadly conserved σ factor that directs transcription of genes involved in flagellar motility. We previously identified FliA-transcribed genes in *Escherichia coli* and *Salmonella enterica* serovar Typhimurium, and we showed that *E. coli* FliA transcribes many unstable, non-coding RNAs from intragenic promoters. Here, we show that FliA in *S*. Typhimurium also directs transcription of large numbers of unstable, non-coding RNAs from intragenic promoters, and we identify two previously unreported FliA-transcribed protein-coding genes. One of these genes, *sdiA*, encodes a transcription factor that responds to quorum sensing signals produced by other bacteria. We show that FliA-dependent transcription of *sdiA* is required for SdiA activity, highlighting a regulatory link between flagellar motility and intercellular communication.

**IMPORTANCE:** Initiation of bacterial transcription requires association of a σ factor with the core RNA polymerase to facilitate sequence-specific recognition of promoter elements. FliA is a widely conserved σ factor that directs transcription of genes involved in flagellar motility. We previously showed that *Escherichia coli* FliA transcribes many unstable, non-coding RNAs from promoters within genes. Here, we demonstrate the same phenomenon in *Salmonella* Typhimurium. We also show that *S*. Typhimurium FliA directs transcription of the *sdiA* gene, which encodes a transcription factor that responds to quorum sensing signals produced by other bacteria. FliA-dependent transcription of *sdiA* is required for transcriptional control of SdiA target genes, highlighting a regulatory link between flagellar motility and intercellular communication.

## INTRODUCTION

In *Escherichia coli* and *Salmonella enterica*, genes involved in flagellar motility are classified into three groups. Class I flagellar genes are *flhD* and *flhC*, which encode the master flagellar regulator, FlhDC. Class II flagellar genes are directly activated by FlhDC and include *fliA*, which encodes FliA/σ^28^, a broadly conserved alternative σ factor. Class III flagellar genes are transcribed by RNA polymerase associated with FliA (RNAP:FliA). The majority of Class III regulation is conserved across bacterial species, with a small degree of species-specific regulation (Fitzgerald *et al.*, 2018).

We previously used a combination of ChIP-seq and RNA-seq to identify sites of RNAP:FliA binding in *E. coli* and *S. enterica* serovar Typhimurium. Thus, we identified novel FliA-transcribed genes, and we showed that in *E. coli*, most FliA-transcribed RNAs initiate within genes, and are likely non-coding and unstable (Fitzgerald *et al.*, 2014; Fitzgerald *et al.*, 2018). These intragenic *E. coli* FliA promoters are not conserved at the sequence level, and we did not detect FliA binding to homologous regions in *S*. Typhimurium. Indeed, we identified few intragenic FliA sites in *S*. Typhimurium. However, we identified fewer FliA sites overall in *S*. Typhimurium than in *E. coli*, suggesting that the *S*. Typhimurium data had reduced sensitivity.

Several *E. coli* and *S*. Typhimurium genes in the flagellar regulatory cascade encode regulators, thereby extending the indirect flagellar regulatory network. Specifically, *flgM* encodes an anti-σ factor that sequesters FliA (Ohnishi *et al.*, 1992); *fliZ* encodes a DNA-binding transcription factor with a large regulon (Pesavento and Hengge, 2012); *yhjH* and *ygcR* encode modulators of c-di-GMP signaling (Boehm *et al.*, 2010; Paul *et al.*, 2010); and *E. coli* FliA transcribes two stable non-coding RNAs (Fitzgerald *et al.*, 2014), one of which, UhpU, has been reported to base pair with multiple mRNAs that may be regulatory targets (Melamed *et al.*, 2016; Melamed *et al.*, 2020).

Here, we show that *S*. Typhimurium RNAP:FliA binds many promoters within genes, likely promoting transcription of unstable, non-coding RNAs, as we have previously described for *E. coli* FliA. Two FliA-transcribed protein-coding genes were also identified. One is a homologue of a known FliA-transcribed gene from *E. coli*. The other, *sdiA*, encodes a transcription factor that responds to quorum sensing signals generated by other bacterial species. Thus, we identified an additional level of regulatory crosstalk that involves FliA, connecting *S*. Typhimurium motility with the sensing of other bacterial species in the gut.

## RESULTS

### Deletion of strong FliA binding sites facilitates identification of weaker sites

We previously identified 23 FliA binding sites in *S*. Typhimurium strain 14028s using ChIP-seq (Fitzgerald *et al.*, 2018), and identified FliA-dependent genes by microarray analysis and RNA-seq (Frye *et al.*, 2006; Fitzgerald *et al.*, 2018). While many FliA binding sites were identified, we reasoned that weaker sites may have been missed. To facilitate identification of these weaker sites, ChIP-seq analysis was performed using N-terminally FLAG_3_-tagged FliA in a “motility promoter-deficient” *S*. Typhimurium LT2 strain that lacks 14 of the known FliA promoters. By this method, 265 FliA-bound regions were identified (Table S1); these include 7 of the regions identified in our earlier study (Fitzgerald *et al.*, 2018). Examples of FliA-bound regions identified by ChIP-seq in the motility promoter-deficient strain are shown in Figure 1A. A strongly enriched sequence motif was identified from 255 of the 265 FliA-bound regions (Figure 1B; Table S1); this motif is an excellent match to the well-described FliA −10 element (Park *et al.*, 2001; Fitzgerald *et al.*, 2014; Fitzgerald *et al.*, 2018); as a group, the FliA-bound regions lack good matches to the FliA −35 element (Figure 1B). Instances of the motif were significantly centrally enriched relative to the ChIP-seq peak centers (Centrimo *E*-value = 8.4e^−12^). We conclude that most, if not all, of the 265 FliA-bound regions identified by ChIP-seq in the motility promoter-deficient strain are genuine.

**Figure 1.**
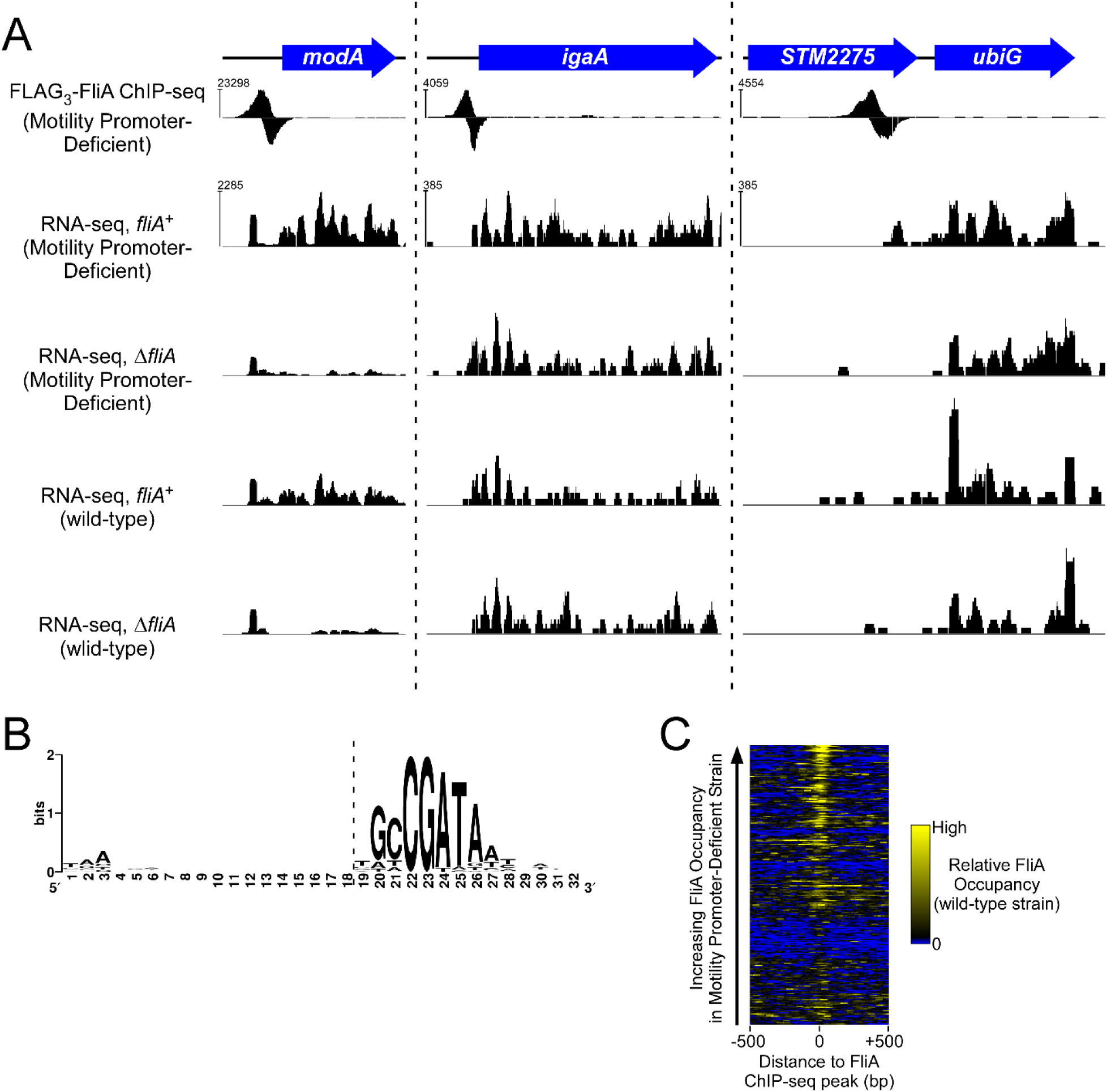
ChIP-seq of FliA in a motility promoter-deficient strain identifies novel binding sites. **(A)** Examples of FliA binding sites identified by ChIP-seq in a motility promoter-deficient strain. The schematic shows the position of genes in three genomic regions. FLAG_3_-FliA (JP095) ChIP-seq sequence read coverage (for one of two replicates) is shown, with positive values indicating coverage on the plus strand and negative values indicating coverage on the minus strand. RNA-seq data are shown for *fliA*+ (LT2 and TH15903) and Δ*fliA* (TH7410 and TH22673) motility promoter-deficient and wild-type strains. The scale on the *y*-axis differs for the ChIP-seq and RNA-seq data, and differs between each of the three panels. The scale on the *y*-axis for RNA-seq data is the same for the four datasets shown within each panel. The values indicated to the left of each panel are the highest read depth values for ChIP-seq or RNA-seq data on that strand in the genomic range shown. **(B)** Extended enriched sequence motif identified in 255 of the 265 FliA-bound regions identified by ChIP-seq using MEME (Bailey and Elkan, 1994). The *y*-axis represents the information content at each position of the motif, and the relative height of each letter indicates the relative proportion of the corresponding base at each position. Positions 1-19 (to the left of the dashed, vertical line) are not part of the motif identified by MEME. **(C)** Heat-map showing sequence read coverage in previously described ChIP-seq data for FLAG_3_-FliA in *S*. Typhimurium 14028s (Fitzgerald *et al.*, 2018). Sequence read coverage, summed for both strands and across two replicates, is shown for 1 kb regions surrounding each of the 255 FLAG_3_-FliA ChIP-seq peak centers identified from the motility promoter-deficient strain, with regions ranked by the level of FLAG_3_-FliA ChIP-seq signal (“occupancy”) in the motility promoter-deficient strain. Yellow indicates high sequence read coverage and blue indicates zero coverage.

To determine whether the novel FliA-bound regions in the motility promoter-deficient strain are also bound by FliA in a wild-type strain, FliA ChIP-seq signals from our previous study were examined (Fitzgerald *et al.*, 2018) at the 259 novel FliA-bound regions identified in the current study. A striking local enrichment of FliA binding at many of the novel sites was identified, especially those that showed higher ChIP-seq signal in the motility promoter-deficient strain (Figure 1C). We conclude that many, perhaps most, of the novel FliA sites are bound in a wild-type strain, but failed to meet the threshold for detection in our previous study.

### Weak, intragenic FliA-dependent promoters are widespread in S. Typhimurium

Our previous study of *E. coli* FliA identified a large number of intragenic FliA-dependent promoters that likely generate unstable, non-coding RNAs (Fitzgerald *et al.*, 2014). To determine if a similar phenomenon occurs in *S*. Typhimurium, the locations of FliA binding sites relative to annotated genes were determined. Of the 255 FliA-bound regions that have an identified FliA binding site, 37 (15.2%) are intergenic (Figure 2A; Table S1). This suggests a random distribution of most FliA binding sites, since 13.5% of the genome is intergenic. To test whether novel FliA binding sites identified in the motility promoter-deficient strain represent active FliA-dependent promoters in a wild-type strain, transcriptional reporter fusions of 10 putative FliA-dependent promoters to *lacZ* were constructed. Specifically, we fused the region from −200 to +10 bp relative to the predicted transcription start site for 10 binding sites that covered a wide range of ChIP-seq occupancy scores, and were not associated with any known or detectable regulation of annotated genes by RNA-seq (Fitzgerald *et al.*, 2018). Expression of the 10 reporter fusions was then measured in *fliA*^+^ and Δ*fliA* cells. For 7 of the fusions, including four for intragenic FliA binding sites, significantly lower expression (t test, *p* < 0.05) was observed in Δ*fliA* cells than in *fliA*^+^ cells (Figure 2B), strongly suggesting that these sites represent active FliA-dependent promoters. Surprisingly the absolute expression level did not correlate with relative ChIP-seq sequence read coverage, suggesting that the dwell time of FliA-containing holoenzymes at different promoters is not strongly correlated with their transcriptional output. An alternative explanation is that expression from plasmid reporters can differ markedly from expression at native chromosomal loci (Bryant *et al.*, 2014).

**Figure 2.**
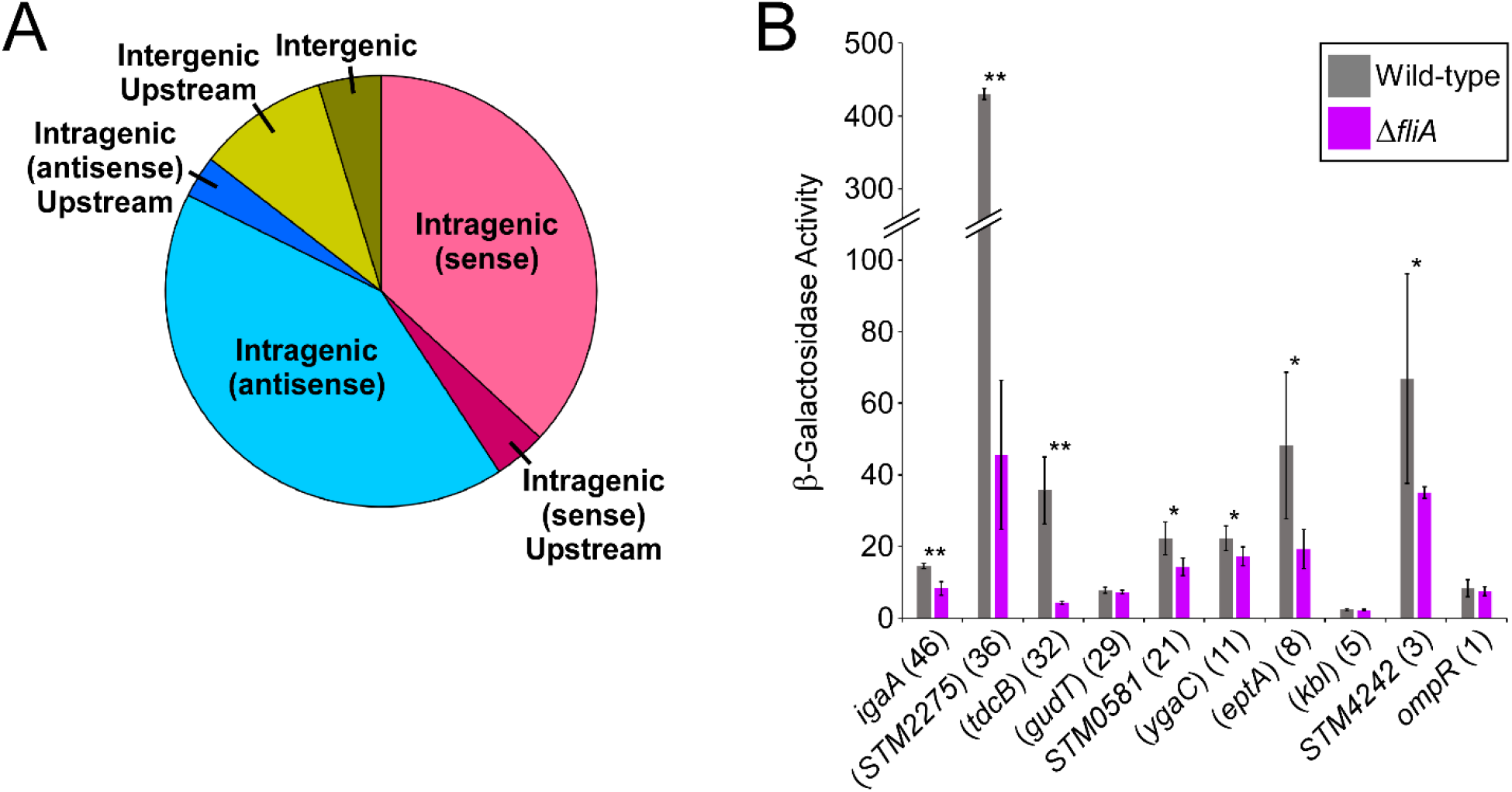
Many novel FliA binding sites represent transcriptionally active FliA promoters inside genes. **(A)** Pie-chart showing the distribution of the 255 FliA binding sites with an associated binding site, relative to annotated genes. “Sense” and “Antisense” refer to the orientation of the predicted FliA promoter relative to the orientation of the overlapping gene. “Upstream” refers to predicted FliA promoters with a predicted promoter that is ≤300 bp upstream of the start of an annotated gene in the same orientation. **(B)** β-galactosidase activity of wild-type (LT2; gray bars) or Δ*fliA* (TH7410; purple bars) cells containing transcriptional fusions of 200 bp regions upstream of ten putative FliA promoters (plasmids pJRP54-72). Gene names not in parentheses indicate intergenic promoters, with the gene being the first downstream gene in the same orientation as the putative promoter. Gene names in parentheses indicate intragenic promoters, with the gene listed being that which encompasses the putative promoter. Numbers in parentheses indicate the relative ChIP-seq sequence read coverage (occupancy scores; see Methods for details) for each putative promoter. Asterisks indicate statistical significance: ** *p* < 0.01, * *p* < 0.05 (one-tailed t test).

### *The* sdiA *and* STM1300 loci *are FliA-transcribed protein-coding genes*

The vast majority of FliA binding sites we identified by ChIP-seq are within or upstream of genes that were not significantly differentially expressed between Δ*fliA* and *fliA*^+^ cells in *S*. Typhimurium 14028s, as determined by RNA-seq (Fitzgerald *et al.*, 2018). To test more rigorously whether these genes are associated with detectable FliA-dependent expression, RNA-seq was used to compare RNA levels between *fliA*^+^ and Δ*fliA* derivatives of motility promoter-deficient and motility-proficient *S*. Typhimurium LT2. The majority of genes with an upstream or internal FliA binding site (Table S1) were not differentially expressed between the *fliA*^+^ and Δ*fliA* strains in either genetic background (*q*-value ≤ 0.01, ≥ 2 higher expression in *fliA*^+^ cells than in Δ*fliA* cells; Table S2), even in cases where we observed FliA-dependent expression with transcriptional reporter fusions (Figure 2B). Thus, the RNA-seq data are consistent with most FliA promoters driving transcription of unstable RNAs that are not detectable by conventional RNA-seq. Nonetheless, we did observe differential expression between *fliA^+^* and Δ*fliA* strains in both genetic backgrounds of six genes with upstream FliA binding sites. As expected, we observed differential expression of *modA* and *STM0567*, as previously described for the equivalent genes in *S*. Typhimurium 14028s (Figure 1A; note that *STM14_0662* is the homologous gene for *STM0567* in strain 14028s) (Fitzgerald *et al.*, 2018). In addition, we observed differential expression of *ybeM*, *ycgO*, STM1300, and *sdiA*, suggesting that mRNAs for these protein-coding genes are transcribed by FliA (Figure 3A). In a previous RNA-seq study, we observed significantly lower expression of *sdiA* and *STM1300* homologues, but not *ybeM* or *ycgO*, in Δ*fliA S*. Typhimurium 14028s than in wild-type cells (Fitzgerald *et al.*, 2018). It is also noteworthy that the putative FliA promoter upstream of *ycgO* was predicted in our earlier study by sequence-based phylogenetic analysis, due to the presence of a conserved FliA binding site upstream of the *E. coli ycgO* homologue (also known as *cvrA*; encodes a putative K^+^:H^+^ antiporter) (Fitzgerald *et al.*, 2014). Expression of *E. coli ycgO* was not found to be significantly different between a wild-type and Δ*fliA* mutant strain (Fitzgerald *et al.*, 2014).

**Figure 3.**
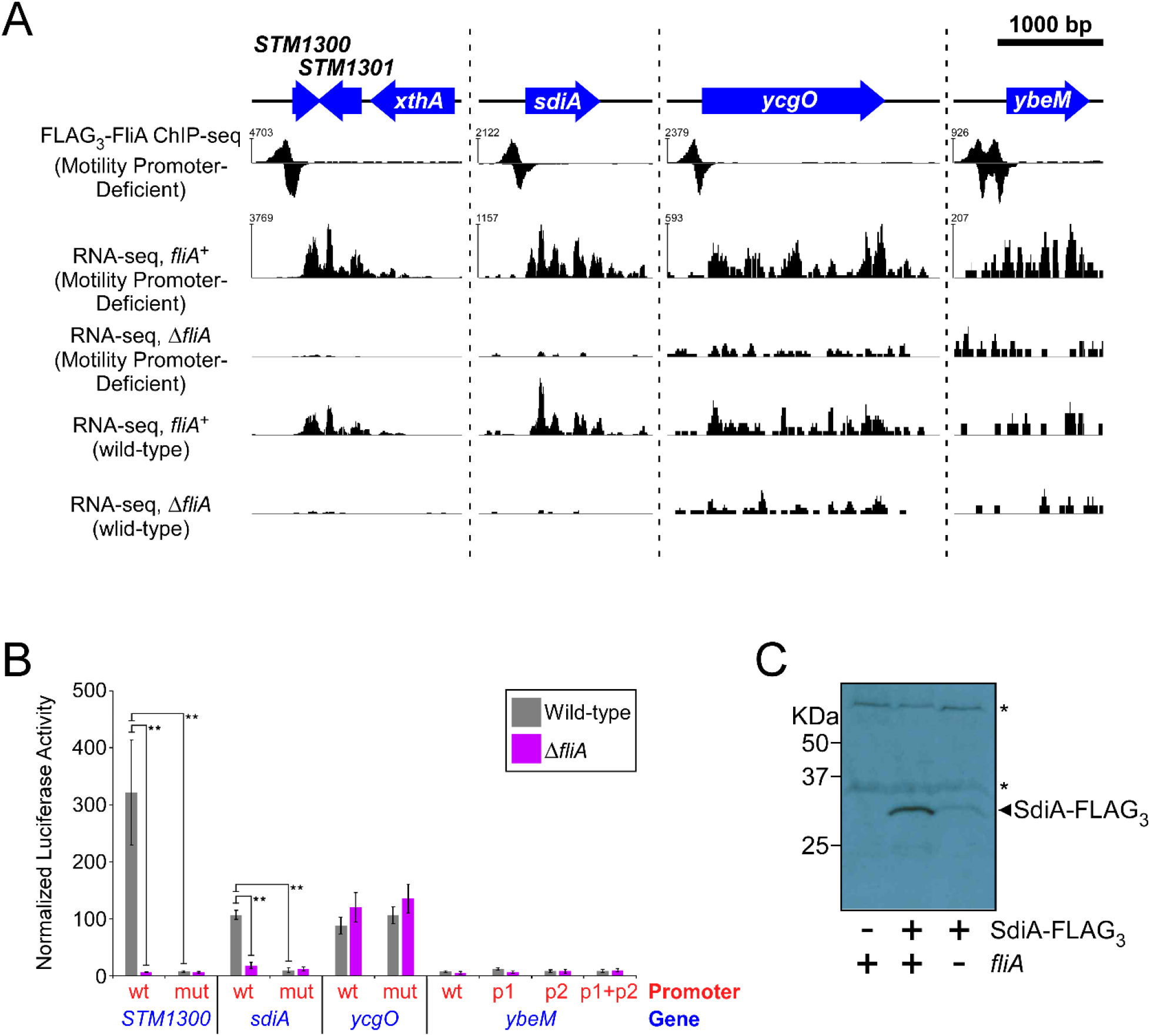
FliA-dependent transcription of the *STM1300* and *sdiA* protein-coding genes. **(A)** FliA binding sites upstream of *STM1300*, *sdiA*, and *ybeM*. The schematic shows the position of genes in the three genomic regions. FLAG_3_-FliA ChIP-seq sequence read coverage (for one of two replicates) is shown, with positive values indicating coverage on the plus strand and negative values indicating coverage on the minus strand. RNA-seq data are shown for *fliA*+ and Δ*fliA* motility promoter-deficient and wild-type strains. The scale on the *y*-axis differs for the ChIP-seq and RNA-seq data, and differs between each of the three panels. The scale on the *y*-axis for RNA-seq data is the same for the four datasets shown within each panel. The values indicated to the left of each panel are the highest read depth values for ChIP-seq or RNA-seq data on that strand in the genomic range shown. **(B)** Luciferase activity of wild-type (LT2) or Δ*fliA* (TH7410) *S*. Typhimurium containing translational reporter fusions for *STM1300*, *sdiA*, *ycgO* or *ybeM* (plasmids pJRP33-34, pJRP37-40, and pJRP43-46). The putative FliA-dependent promoters are wild-type or were mutated to prevent RNAP:FliA binding. In the case of *ybeM*, there are two predicted FliA-dependent promoters, and mutations were made in each individually (p1, p2), or both simultaneously (p1 + p2). Statistically significant differences (t-test *p* < 0.01) between wild-type *fliA* and Δ*fliA* strains or between wild-type and mutant reporter fusions are indicated by a double asterisk. **(C)** Western blot showing SdiA-FLAG_3_ in wild-type (LT2; untagged control), SdiA-FLAG_3_ (JP085), or SdiA-FLAG_3_ Δ*fliA* (JP089) *S*. Typhimurium. The expected position of SdiA-FLAG_3_ is indicated. Asterisks indicate non-specific bands.

To test more directly whether *ybeM*, *ycgO*, *STM1300*, and *sdiA* are transcribed by FliA, translational reporter fusions to luciferase were constructed with the upstream regions, including the putative promoters, up to the annotated start codons. Equivalent reporter fusions with point mutations in the putative −10 element for FliA were constructed; these mutations were expected to prevent transcription by RNAP:FliA. In the case of *ybeM*, three different promoter mutations were made because the ChIP-seq data identified two distinct FliA binding sites, 114 bp apart from one another. Specifically, reporter fusion strains were constructed with either individual promoter mutations, or with mutations in both promoters. The reporter constructs were introduced into wild-type (*fliA*^+^) *S*. Typhimurium LT2 or a Δ*fliA* derivative, and luciferase activity was measured (Figure 3B). Robust luciferase activity was observed with reporter fusions to both *sdiA* and *STM1300* upstream regions, with activity dependent upon an intact promoter and the presence of *fliA*. By contrast, expression of the *ycgO* reporter fusion was unchanged by mutation of the putative promoter, and expression of the wild-type reporter was similar in *fliA*^+^ and Δ*fliA* strains. Very weak expression was detected for the reporter fusion to *ybeM*, and expression was unchanged by mutation of either or both of the putative promoters, or deletion of *fliA*. We conclude that *sdiA* and *STM1300* are *bona fide* FliA-transcribed genes, whereas *ybeM* is likely not FliA-transcribed. Alternatively, the *ybeM* start codon could be misannotated, which would explain the very weak observed expression. *ycgO* is also likely not FliA-transcribed, although FliA-dependent expression could be masked by substantially higher levels of transcription involving a different Sigma factor.

The *sdiA* gene encodes a DNA-binding transcription factor that senses acyl homoserine lactone (AHL) molecules produced as quorum sensing signals that facilitate gene regulation as a function of cell density (Ahmer, 2004; Waters and Bassler, 2005). *S*. Typhimurium does not encode an AHL synthase (Ahmer, 2004), but SdiA has been shown to respond to AHLs produced by other bacteria (Smith and Ahmer, 2003), including within mammalian and reptilian hosts (Smith *et al.*, 2008; Dyszel *et al.*, 2010). In the presence of AHLs, SdiA activates transcription of several genes, including *rck*, which encodes an outer membrane invasin (Abed *et al*., 2014). To determine the impact of FliA on SdiA protein levels under a more relevant growth condition, we tagged the chromosomal copy of *sdiA* with three FLAG epitopes and measured SdiA-FLAG_3_ protein levels by Western blot in wild-type or Δ*fliA* cells grown to high density in the presence of the AHL 3OC_6_HSL, a condition known to induce SdiA activity (Smith and Ahmer, 2003); an untagged wild-type strain served as a negative control (Figure 3C). SdiA-FLAG_3_ protein levels were considerably higher in wild-type cells than in Δ*fliA* cells (Figure 3C).

### FliA is required for activity of the SdiA transcription factor

We hypothesized that FliA-dependent transcription of *sdiA* is required for *S*. Typhimurium to accumulate sufficient SdiA protein to activate transcription of its target genes. To test this hypothesis, expression of an *rck* reporter fusion to luciferase was assayed in wild-type, Δ*sdiA*, and Δ*fliA* cells. We measured luciferase activity in cells grown to high density in the presence or absence of 3OC_6_HSL (Figure 4A). Consistent with a previous study (Smith and Ahmer, 2003), increased expression of the reporter fusion was observed for wild-type cells grown with 3OC_6_HSL, but not for wild-type cells grown without 3OC_6_HSL, nor for Δ*sdiA* cells grown with or without 3OC_6_HSL. This result supports the established function of SdiA as an AHL-responsive transcriptional activator of *rck*. Strikingly, expression of the reporter fusion in the Δ*fliA* strain was similar to that in the Δ*sdiA* strain, strongly suggesting that levels of SdiA in Δ*fliA* cells are insufficient to activate transcription of *rck*.

**Figure 4.**
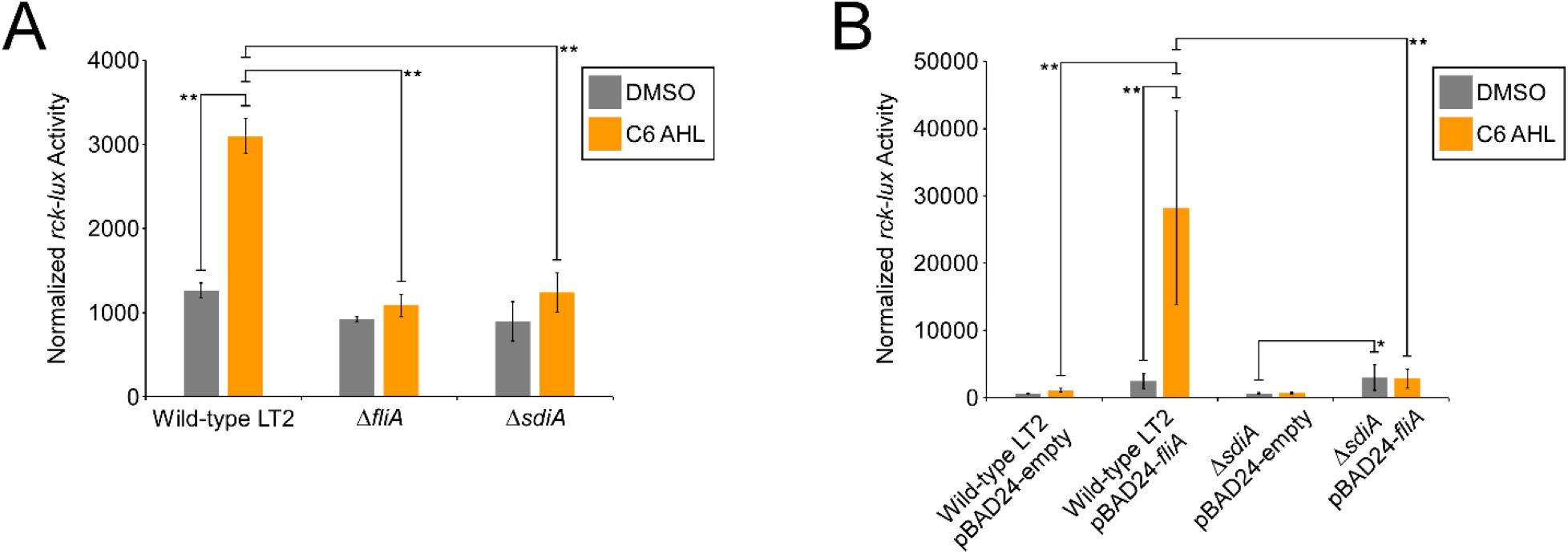
*fliA* is required for cells to accumulate normal levels of active SdiA. **(A)** Luciferase activity of wild-type (LT2), Δ*fliA* (JP096), or Δ*sdiA* (JP087) *S*. Typhimurium containing a reporter fusion for *rck* (pBA428). Cells were grown with either DMSO or 3OC_6_HSL, as indicated. **(B)** Luciferase activity of wild-type (LT2) or Δ*sdiA* (JP087) *S*. Typhimurium containing a reporter fusion for *rck* (pBA428) and either empty pBAD24 or pBAD24 expressing *fliA* (pJRP81). Cells were grown with either DMSO or 3OC_6_HSL, as indicated. Statistically significant differences are indicated by a single asterisk (t-test *p* < 0.05) or a double asterisk (t-test *p* < 0.01).

To test whether the effect of deleting *fliA* on expression of *rck* is due a reduction in the level of SdiA, expression of the *rck*-luciferase reporter fusion was measured in wild-type or Δ*sdiA* cells that contained a plasmid overexpressing *fliA* or an equivalent empty vector (Figure 4B). Overexpression of *fliA* in wild-type cells led to a large increase in expression of the reporter fusion for cells grown with 3OC_6_HSL, but not for cells grown without 3OC_6_HSL. The same increase in expression of the reporter was not observed in Δ*sdiA* cells overexpressing *fliA*. We conclude that FliA-dependent transcription of *sdiA* is required for *S*. Typhimurium to accumulate sufficient SdiA protein to activate transcription of its target genes.

Interestingly, overexpression of *fliA* resulted in a modest, statistically significant (t-test *p* < 0.05) increase in *rck* expression in both wild-type and Δ*sdiA* cells grown without 3OC_6_HSL, and in Δ*sdiA* cells grown with 3OC_6_HSL. These data suggest that there is a small effect of FliA on *rck* expression that is independent of SdiA.

## DISCUSSION

### Numerous weak, intragenic FliA promoters in S. Typhimurium

Our previous studies identified 52 FliA promoters in *E. coli*, of which most are intragenic, likely driving transcription of unstable, non-coding RNAs (Fitzgerald *et al.*, 2014; Fitzgerald *et al.*, 2018). Moreover, very few of these intragenic promoters are conserved in other species (Fitzgerald *et al.*, 2018), suggesting they arise due to genetic drift. Here, we show that there are many analogous FliA promoters in *S*. Typhimurium. These promoters tend to bind RNAP:FliA with lower affinity, as evidenced by our failure to detect most of them using ChIP-seq in a wild-type strain (Fitzgerald *et al.*, 2018). Consistent with lower affinity for FliA, these promoters tend to rely on the −10 element, and have more degenerate −35 elements than the canonical, intergenic FliA promoters described previously (Frye *et al.*, 2006; Fitzgerald *et al.*, 2018). Although we identified considerably more intragenic FliA promoters in *S*. Typhimurium than we did in *E. coli*, we speculate that this is due to the increased sensitivity achieved by using a motility promoter-deficient strain of *S*. Typhimurium. If, as we have hypothesized, intragenic FliA promoters arise due to genetic drift, we would expect there to be a continuum of binding of RNAP:FliA to sites across the chromosome. Hence, a more sensitive assay for detecting FliA-DNA interactions would inevitable identify more FliA promoters. In other words, we propose that there is not a set of sequences that constitute FliA promoters and another set that does not; rather all sequences can serve as FliA promoters across a very large range of affinities.

### STM1300 *and* sdiA *are novel FliA-transcribed protein-coding genes*

Although our data suggest that most of the novel FliA promoters are non-functional, and drive transcription of non-coding RNAs, we identified two FliA-transcribed protein-coding genes: *sdiA* and *STM1300*. While these genes have not been established in prior work to be FliA-transcribed in *S*. Typhimurium, a previous study showed that expression of each of *sdiA* and *STM1300* correlates with that of Class II and Class III flagellar genes, respectively (Wang *et al.*, 2006). Our previous study of *E. coli* FliA strongly suggests that FliA-dependent transcription of *sdiA* is not conserved in *E. coli* (Fitzgerald *et al.*, 2014). By contrast, *STM1300* has a close homologue in *E. coli*, *ynjH* (73% amino acid identity, 99% coverage), which we and others have previously reported to be transcribed by FliA (Yu *et al.*, 2006; Zhao *et al.*, 2007; Fitzgerald *et al.*, 2014). Thus, *STM1300*/*ynjH* is a conserved member of the FliA regulon. There is currently no known function for *STM1300*/*ynjH*.

### *FliA-dependent transcription of* sdiA *may coordinate motility with interbacterial communication*

*Salmonella* is unusual in that it expresses a LuxR homologue, SdiA, but does not produce AHLs. Hence, SdiA’s primary function is likely to be sensing of AHLs produced by other bacteria. Consistent with this, SdiA activity, which is AHL-dependent, has been demonstrated in *S*. Typhimurium cells growing within mammalian and reptilian hosts (Smith *et al.*, 2008; Dyszel *et al.*, 2010). The complete SdiA regulon is not known, but seven SdiA-activated genes have been identified (Ahmer *et al.*, 1998; Michael *et al.*, 2001; Smith and Ahmer, 2003). However, these genes are largely uncharacterized, so the physiological significance of SdiA-mediated regulation is unclear. Our data suggest show that expression of SdiA-dependent genes is tightly coupled to expression of flagellar genes. Thus, SdiA may be involved in coordinating the response to surrounding bacteria with motility.

We extracted the sequences upstream of *sdiA* homologues in species from 65 different γ-proteobacterial genera and identified the best match within these sequences to the known *E. coli* FliA binding site motif (Fitzgerald *et al.*, 2014). Thus, we can predict the degree to which the FliA promoter upstream of *sdiA* is conserved in other species. While the FliA binding site upstream of *sdiA* is not broadly conserved, we identified species of *Enterobacter*, *Citrobacter*, *Kosakonia*, and *Cronobacter* with better-scoring sequences than the site from *S*. Typhimurium LT2 (Table S3). We conclude that FliA-dependent transcription of *sdiA* is likely conserved in a small number of closely related genera. Species of *Enterobacter* and *Citrobacter* have been reported to produce AHLs, including 3OC_6_HSL (Lau *et al.*, 2013; Kher *et al.*, 2019), and at least one species of *Enterobacter* that is known to produce 3OC_6_HSL has a high-scoring potential FliA binding site upstream of *sdiA* (Lau *et al.*, 2013; Lau *et al.*, 2014). Thus, it is likely that, in some species, FliA contributes to the response to endogenously produced AHLs.

## MATERIALS AND METHODS

### Strains

All strains used in this work are derivatives of *S*. Typhimurium LT2, and are listed in Table 1. All oligonucleotides used in this work are listed in Table S4. Strains TH15903 and TH22673 were made by successive deletion of σ^28^-dependent genes and associated promoter regions. The details of construction of these strains are provided in the Supplementary Methods. Strain TH7410 (LT2 Δ*fliA5805*::*tetRA*) has been described previously (Frye *et al.*, 2006). Strains JP085 (LT2 *sdiA*-FLAG_3_), JP096 (LT2 Δ*fliA*), and JP087 (LT2 Δ*sdiA*) are derivatives of LT2 that were constructed using the FRUIT recombineering method (Stringer *et al.*, 2012) with oligonucleotides JW10058+9 (JP085), JW6298-303 (JP096), and JW10106-111 (JP087). JP095 (Δ*flgMN7753* Δ*flgKL7770* Δ*trg-7774* Δ*ycgR7775* Δ*mcpA7792* Δ*fliB-T7771* Δ*tsr-7828* Δ*mcpB7835* Δ*hin-fljA7752* Δ*prgH-hilA7791* Δ*cheV7829* Δ*tcp-7827* Δ*yhjH7740* Δ*aer-mcpC7834* P*motA7795* Δ*motA-cheZ7888* FLAG_3_-*fliA*) is a derivative of TH15903 that was constructed using FRUIT with oligonucleotides JW6306+7. JP089 (LT2 Δ*fliA5805*::*tetRA sdiA*-FLAG_3_) is a derivative of TH7410 that was constructed using FRUIT with oligonucleotides JW10058+9.

**Table 1.** List of strains used in this study.

| Name | Description | Source* |
| --- | --- | --- |
| LT2 | Wild type | John Roth |
| TH7410 | $\Delta fliA5805::tetRA$ | (Frye *et al.*, 2006) |
| TH14224 | $\Delta flgKL7719::tetRA$ | |
| TH14226 | $\Delta tsr-7720::tetRA$ | |
| TH14228 | $\Delta tcp-7721::tetRA$ | |
| TH14230 | $\Delta trg-7722::tetRA$ | |
| TH14232 | $\Delta mcpB7723::tetRA$ | |
| TH14234 | $\Delta aer-mcpC7724::tetRA$ | |
| TH14236 | $\Delta cheV7725::tetRA$ | |
| TH14237 | $\Delta prgH-hilA7726::tetRA$ | |
| TH14238 | $\Delta fliB-T7727::tetRA$ | |
| TH14239 | $\Delta yhjH7728::tetRA$ | |
| TH14301 | $\Delta hin-fljA7731::tetRA$ | |
| TH14303 | $\Delta flgMN7732::tetRA$ | |
| TH14305 | $\Delta mcpA7733::tetRA$ | |
| TH14554 | $\Delta yhjH7740$ | |
| TH14589 | $\Delta ycgR7751::tetRA$ | |
| TH14591 | $\Delta hin-fljA7752$ | |
| TH14592 | $\Delta flgMN7753$ | |
| TH14690 | $\Delta flgKL7770$ | |
| TH14691 | $\Delta fliB-T7771$ | |
| TH14735 | Δtrg-7774 | |
| TH14779 | ΔycgR7775 | |
| TH14864 | ΔprgH-hilA7791 | |
| TH14868 | ΔmcpA7792 | |
| TH15143 | ΔmotA-cheZ7809::tetRA | |
| TH15171 | Δtcp-7827 | |
| TH15172 | Δtsr-7828 | |
| TH15173 | ΔcheV7829 | |
| TH15356 | Δaer-mcpC7834 | |
| TH15357 | ΔmcpB7835 | |
| TH15901 | ΔmotA-cheZ7888 | |
| TH15903 | ΔflgMN7753 ΔflgKL7770 Δtrg-7774 ΔycgR7775 ΔmcpA7792 ΔfliB-T7771 Δtsr-7828 ΔmcpB7835 Δhin-fljA7752 ΔprgH-hilA7791 ΔcheV7829 Δtcp-7827 ΔyhjH7740 Δaer-mcpC7834 P motA7795 ΔmotA-cheZ7888 | |
| TH22673 | ΔflgMN7753 ΔflgKL7770 Δtrg-7774 ΔycgR7775 ΔmcpA7792 ΔfliB-T7771 Δtsr-7828 ΔmcpB7835 Δhin-fljA7752 ΔprgH-hilA7791 ΔcheV7829 Δtcp-7827 ΔyhjH7740 Δaer-mcpC7834 P motA7795 ΔmotA-cheZ7888 ΔfliA5805::tetRA | |
| JP085 | LT2 sdiA-FLAG<sub>3</sub> |  |
| JP096 | LT2 ΔfliA | |
| JP087 | LT2 ΔsdiA | |
| JP095 | ΔflgMN7753 ΔflgKL7770 Δtrg-7774 ΔycgR7775 ΔmcpA7792 ΔfliB-T7771 Δtsr-7828 ΔmcpB7835 Δhin-fljA7752 ΔprgH-hilA7791 ΔcheV7829 Δtcp-7827 ΔyhjH7740 Δaer-mcpC7834 P motA7795 ΔmotA-cheZ7888 FLAG<sub>3</sub>-fliA | |
| JP089 | LT2 ΔfliA5805::tetRA sdiA-FLAG<sub>3</sub> | |

\*Unless indicated otherwise, all strains were constructed for this work

### Plasmids

All plasmids are listed in Table 2. All *lacZ* transcriptional reporter fusions were constructed in plasmid pAMD-BA-*lacZ* (Stringer *et al.*, 2014). 10 putative promoter regions, from −200 to +10 bp relative to the predicted transcription start site (predicted based on the position of the putative FliA binding site), were PCR-amplified from *S*. Typhimurium LT2 genomic DNA using oligonucleotides listed in Table S4. PCR products were cloned using the In-Fusion method (Clontech) into pAMD-BA-*lacZ* digested with *Nhe*I and *Sph*I to generate plasmids pJRP54-72 (see Table 2 for details).

**Table 2.** List of plasmids used in this study.

| Name | Description | Source* |
| --- | --- | --- |
| pKD46 | Para- λRed recombinase (Rep<sup>ts</sup> Ap<sup>R</sup>) | (Datsenko and Wanner, 2000) |
| pBA428 | rck transcriptional luciferase fusion | (Michael *et al.*, 2001) |
| pGB135 | Empty lucCDABE (luciferase) reporter plasmid | (Baniulyte and Wade, 2019) |
| pJRP33 | pGB135 translational luciferase fusion, STM1300 upstream region |  |
| pJRP34 | pGB135 translational luciferase fusion, *STM1300* upstream region, putative FliA site mutated |  |
| pJRP37 | pGB135 translational luciferase fusion, *sdiA* upstream region |  |
| pJRP38 | pGB135 translational luciferase fusion, *sdiA* upstream region, putative FliA site mutated |  |
| pJRP39 | pGB135 translational luciferase fusion, *ycgO* upstream region |  |
| pJRP40 | pGB135 translational luciferase fusion, *ycgO* upstream region, putative FliA site mutated |  |
| pJRP43 | pGB135 translational luciferase fusion, *ybeM* upstream region |  |
| pJRP44 | pGB135 translational luciferase fusion, *ybeM* upstream region, putative FliA site #1 mutated |  |
| pJRP45 | pGB135 translational luciferase fusion, *ybeM* upstream region, putative FliA site #1 mutated |  |
| pJRP46 | pGB135 translational luciferase fusion, *ybeM* upstream region, putative FliA sites #1 and #2 mutated |  |
| pAMD-BA-*lacZ* | Empty *lacZ* reporter plasmid | (Stringer *et al.*, 2014) |
| pJRP54 | pAMD-BA-*lacZ* transcriptional *lacZ* fusion, putative FliA-dependent promoter upstream of *igaA* |  |
| pJRP56 | pAMD-BA-*lacZ* transcriptional *lacZ* fusion, putative FliA-dependent promoter within *STM2275* |  |
| pJRP58 | pAMD-BA-*lacZ* transcriptional *lacZ* fusion, putative FliA-dependent promoter within *tdcB* |  |
| pJRP60 | pAMD-BA-*lacZ* transcriptional *lacZ* fusion, putative FliA-dependent promoter within *gudT* |  |
| pJRP62 | pAMD-BA-*lacZ* transcriptional *lacZ* fusion, putative FliA-dependent promoter upstream of *STM0581* |  |
| pJRP64 | pAMD-BA-*lacZ* transcriptional *lacZ* fusion, putative FliA-dependent promoter within *ygaC* |  |
| pJRP66 | pAMD-BA-*lacZ* transcriptional *lacZ* fusion, putative FliA-dependent promoter within *eptA* |  |
| pJRP68 | pAMD-BA-*lacZ* transcriptional *lacZ* fusion, putative FliA-dependent promoter within *kbl* |  |
| pJRP70 | pAMD-BA-*lacZ* transcriptional *lacZ* fusion, putative FliA-dependent promoter upstream of *STM4242* |  |
| pJRP72 | pAMD-BA-*lacZ* transcriptional *lacZ* fusion, putative FliA-dependent promoter upstream of *ompR* |  |
| pBAD24 | Empty plasmid with P<sub>BAD</sub> promoter for arabinose-inducible expression | (Guzman *et al.*, 1995) |
| pJRP81 | pBAD24-*fliA* |  |

\*Unless indicated otherwise, all plasmids were constructed for this work

All luciferase (*luxCDABE* from *Photorhabdus luminescens*) translational reporter fusions were constructed in plasmid pGB135 (Baniulyte and Wade, 2019). Wild-type and mutant upstream regions, including the putative promoter and extending to the annotated start codon, were PCR-amplified from *S*. Typhimurium LT2 genomic DNA using oligonucleotides listed in Table S4. Mutant promoter PCR products were generated by SOEing PCR (Horton *et al.*, 1990). PCR products were cloned using the In-Fusion method (Clontech) into pGB135 digested with *Bam*HI and *Eco*RI to generate plasmids pJRP33-34, pJRP37-40, and pJRP43-46 (see Table 2 for details).

pBA428 is a transcription reporter fusion of *rck* to luciferase (*luxCDABE* from *P. luminescens*), and has been described previously (Michael *et al.*, 2001).

pJRP81 is a derivative of pBAD24 (Guzman *et al.*, 1995) that contains *fliA*. The *fliA* gene was PCR-amplified from *S*. Typhimurium LT2 genomic DNA using oligonucleotides JW10331+2. The PCR product was cloned using the In-Fusion method (Clontech) into pBAD24 digested with *Kpn*I and *Sph*I to generate pJRP81.

### ChIP-seq

JP095 was grown in LB to an OD_600_ of ~1. ChIP-seq libraries from two independent biological replicates were prepared as previously described (Stringer *et al.*, 2014). Libraries were sequenced on a NextSeq (Illumina) by the Wadsworth Center Applied Genomic Technologies Core Facility. ChIP-seq sequence reads were aligned to the *S*. Typhimurium LT2 chromosome using CLC Genomics Workbench (version 8). ChIP-seq peaks were called using a previously described analysis pipeline (Fitzgerald *et al.*, 2014). Occupancy scores listed in Table S1 are measures of relative sequence read coverage from ChIP-seq data, averaged across two replicates. ChIP-seq peaks were identified only in regions exceeding a threshold sequencing read depth. Occupancy scores indicate the fold difference between the sequence read coverage at the corresponding ChIP-seq peaks and the threshold coverage needed to call a peak.

### FliA Motif Discovery

101 bp sequences surrounding ChIP-seq peaks were extracted (overlapping regions were merged) and analyzed by MEME (version 5.1.0, default parameters) (Bailey and Elkan, 1994; Bailey *et al.*, 2009). The position of the inferred motif relative to ChIP-seq peak centers was analyzed using Centrimo (version 5.1.0, default parameters) (Bailey and Machanick, 2012) through the MEME-ChIP tool (Machanick and Bailey, 2011).

### *RNA-*seq

Four independent biological replicates of strains LT2, TH6827, TH15903 and TH22673 were grown in LB to an OD_600_ of ~0.5 at 37 °C. Total RNA was extracted from these strains using a Qiagen RNA purification kit according to the manufacturer’s protocol. Illumina TruSeq HT stranded mRNA library preparations (without polyA selection) were prepared at the University of Utah High-Throughput Genomics Core Facility. Libraries were sequenced on an Illumina HiSeq 2500 instrument (50-cycle single-read sequencing, version 4). Libraries had between 14.8 and 18.4 million aligned reads, with 97-98% of all reads in each dataset aligning to ribosomal RNA. RNA-seq data were analyzed using Rockhopper (version 2.0.3) (McClure *et al.*, 2013).

### β-galactosidase assays

3 ml cultures were grown in LB + 30 μg/mL chloramphenicol at 37 °C with shaking to an OD_600_ of ~1. β-galactosidase assays were conducted as previously described (Stringer *et al.*, 2014). Arbitrary assay units were calculated as 1,000 * A_420_/((A_600_)*(reaction time)).

### Luciferase assays

For luciferase assays with plasmids pJRP33-46, 5 ml cultures were grown in LB + 25 μg/mL kanamycin at 37 °C with shaking to an OD_600_ of ~1. For luciferase assays with pBA428, 5 ml cultures were grown in LB + 20 μg/mL tetracycline + 1 μM 3OC_6_HSL (in DMSO, which was at 0.01% final concentration) or 0.01% DMSO at 37 °C with shaking for 6 hours. 100 μg/mL ampicillin and 0.2% arabinose were also included in the growth medium for cells containing pBAD24 or pJRP81. After growth, 200 μL cells were aliquoted into a 96-well plate with four technical replicates each. Luminescence readings were taken using a Biotek Synergy 2 instrument. Luminescence counts (RLU) were reported as RLU/OD_600_.

### Western blot

Cultures of LT2, JP085 (LT2 *sdiA*-FLAG_3_), and JP089 (LT2 □ *fliA5805*::*tetRA sdiA*-FLAG_3_) were grown in LB + 1 μM 3OC_6_HSL for 6 hours and pelleted. 500 μL cells were pelleted and resuspended in 25 μL Laemmli sample buffer with a 19:1 ratio of buffer to β-mercaptoethanol. Samples were separated on a 10% acrylamide gel. Proteins were transferred to PVDF membrane and probed with M2 anti-FLAG antibody (Sigma; 1 in 1,000 dilution) and HRP-conjugated goat anti-mouse antibody (1 in 250,000 dilution). Tagged proteins were visualized using the ECL Plus detection kit (Lumigen).

### *Phylogenetic comparison of* sdiA *upstream regions*

We used RSAT (Medina-Rivera *et al.*, 2015) (“get-orthologs” tool) to extract the sequence upstream of *sdiA* in 1,990 species covering 65 genera, using *E. coli sdiA* as a reference sequence. Specifically, we extracted upstream sequence from the nearest upstream gene boundary to the position immediately upstream of *sdiA*. We then scored each 28mer in the sense orientation within each of the 1,990 sequences, using a position-weight matrix (PWM) representing all of the FliA binding sites we previously reported in *E. coli* (Fitzgerald *et al.*, 2014). To calculate the scores, we matched each nucleotide within a 28mer to the corresponding nucleotide within the PWM, and multiplied the corresponding frequencies from the PWM, and multiplied by 4 for each position. We then selected the best score for each of the 1,990 sequences. Lastly, we grouped genomes with identical sequences and arbitrarily selected one genome from each of the 627 groups. The score for each unique sequence and the corresponding genome are indicated in Table S3.

## ACKNOWLEDGEMENTS

We thank the Wadsworth Center Applied Genomic Technologies Core Facility and the University of Utah High-Throughput Genomics Core Facility for DNA sequencing. We thank the Wadsworth Center Tissue Culture and Media Core Facility and Glassware Facility for technical support. We thank Jon Paczkowski for helpful discussions. We thank Brian Ahmer for sharing reagents. This work was supported by National Institutes of Health grant R01GM114812 (JTW) and NIH grant R01GM056141 (KTH). This research was supported in part by an appointment to the Infectious Disease (ID) Laboratory Fellowship Program administered by the Association of Public Health Laboratories (APHL) and funded by the Centers for Disease Control and Prevention (CDC).

## DATA AVAILABILITY

Raw ChIP-seq and RNA-seq data are available at from EBI ArrayExpress using accession numbers E-MTAB-9503 and E-MTAB-9504, respectively.

## SUPPLEMENTARY FILES

Supplementary Methods. Construction of strains TH15903 and TH22673.

Table S1. List of ChIP-seq peaks for FLAG_3_-FliA in the motility promoter-deficient strain.

Table S2. Analysis of RNA-seq data for wild-type LT2, Δ*fliA*, motility promoter-deficient *fliA*^+^, and motility promoter-deficient Δ*fliA* strains.

Table S3. List of *sdiA* upstream regions from different γ-proteobacterial species, with the score for the closest match to the FliA binding site position weight matrix.

Table S4. List of oligonucleotides used in this study.

