## Supplementary Methods for "Regulatory crosstalk between motility and interbacterial communication in *Salmonella* Typhimurium"

*Construction of strains* *TH15903 and TH22673*

The E medium of Vogel and Bonner (1), supplemented with 0.2% D-glucose, was used as minimal medium. Lysogeny broth (LB, 10g/l tryptone, 5 g/l yeast extract and 5 g/l NaCl) was sued as rich medium, and when needed, supplemented with the following antibiotics as follows (final concentrations given): tetracycline hydrochloride (Tc) (15 µg/ml), kanamycin sulfate (Km) (50 µg/ml), chloramphenicol (Cm) (12.5 µg/ml), and sodium ampicillin (100 µg/ml for multicopy plasmid selection). Solid medium was supplemented with 12 g/liter Apex chemical reagents agar. The generalized transducing phage of *S.* Typhimurium P22 *HT105/1 int-201* was used in all transductional crosses (2). P22 transducing lysates are prepared by adding 4 ml of P22 broth (LB + E medium salts + 0.2% glucose + P22 *HT105/1 int-201* at ~10^8^ particles/ml) to 1 ml of an overnight culture of cells and incubated at 37°C for 6 to 9 hours. Cells were pelleted in a table-top centrifuge, and 0.4 ml CHCl_3_ was added to the decanted broth, vortexed to kill remaining bacteria, and stored at 4 °C. For a typical transduction, the phage lysate was diluted 10^-3^ in 0.85% sterile saline, and 0.1 ml of the diluted lysate was added to 0.1 ml of a saturated cell culture directly on selective medium. If phenotypic expression was required prior to selection, the phage/cell mixture was placed at 37 °C for 30 min prior to plating. For transductional crosses selecting for tetracycline sensitivity (Tc^S^) where there was a high background of spontaneous Tc^S^ mutants, 0.1 ml of a 10^-1^ phage dilution was incubated with 0.1 ml cells at 37 °C for 30 min, and separately 0.1 ml of saline is incubated with 0.1 ml cells (cell-only control). The mixtures were then diluted 10^-3^ in saline, and 0.1 ml was plated on Tc^S^ selection plates (12 g/l Apex agar, 5 g/l tryptone, 5 g/l yeast extract, 10 g/l NaCl, 10 g/l NaH_2_PO_4_*H_2_O, 12 mg/l fusaric acid, 1 µM ZnCl_2_, and 0.2 mg/ml anhydrotetracycline). One liter of Tc^S^ selection plates was prepared by separately sterilizing the agar, tryptone, and yeast in one flask and the NaCl and NaH_2_PO_4_*H_2_O in a separate flask. When the autoclaved flasks were cooled to ~55°C, the fusaric acid, ZnCl_2_ and anhydrotetracycline were added from sterile stock solutions, the flasks were mixed and poured. All Tc^S^ transductants were verified by PCR or DNA sequence analysis.

Strains TH15903 and TH22673 were made using a combination of chromosomal targeting by λ-Red mediated recombination (recombineering), as described (Karlinsey, 2007), and P22 transductions. Details of strain construction, including all intermediate strains, are listed, below. Abbreviations used: Cm^R^ = chloramphenicol resistance, Cm^S^ = chloramphenicol sensitivity, Km^R^ = kanamycin resistance, Km^S^ = kanamycin sensitivity, Tc^R^ = tetracycline resistance, Tc^S^ = tetracycline sensitivity, PyrC^+^ = growth on minimal glucose medium without added uracil.

TH14224 (Δ*flgKL7719*::*tetRA*) was constructed from LT2 using recombineering with Tc^R^ selection, with oligonucleotides 3235_FlgKL-tetR and 3236_FlgKL-tetA.

TH14690 (Δ*flgKL7770*) was constructed by removing the *tetRA* cassette from TH1424 using recombineering with Tc^S^ selection, with oligonucleotides 3402-FlgKLdelfwd and 3403_FlgKLdelrev.

TH14767 (Δ*pyrC3051*::FKF Δ*flgM5794*::FCF Δ*flgKL7770*) was constructed by P22 transduction with Km^R^ selection of the Δ*pyrC3051*::FKF-linked Δ*flgM5794*::FCF allele from TH10332 (Δ*pyrC3051*::FKF Δ*flgM5794*::FCF; Hughes lab strain collection) into TH14690. Co-transduction of Δ*flgM5794*::FCF was confirmed by PCR.

TH14592 (Δ*flgMN7753*) was constructed by removing the *tetRA* cassette from Δ*flgMN7732*::*tetRA* using recombineering with Tc^S^ selection, with oligonucleotides 3401_FlgMNdelfwd and 3368_FlgMNdelrev2.

TH14976 (Δ*flgMN7753* Δ*flgKL7770*) was constructed by P22 transduction of the *pyrC*-linked Δ*flgMN7753* allele into TH14767, selecting for PyrC^+^, and screening for Cm^S^.

Additional strains were each made in four steps: (i) replacement of a gene and promoter in LT2 with a *tetRA* cassette, using recombineering with Tc^R^ selection, to generate strain “A”, (ii) P22 transduction with Tc^R^ selection of the *tetRA* cassette from “A” into an earlier strain, to generate strain “B”, (iii) removal of the *tetRA* cassette in strain “A”, using recombineering with Tc^S^ selection, to generate strain “C”, and (iv) P22 transduction with Tc^S^ selection of the marker-free deletion from strain “C” into strain “B”, to generate strain “D”.

Starting strain: TH14976 (Δ*flgMN7753* Δ*flgKL7770*)

A: TH14230 (Δ*trg-7722*::*tetRA*); made using oligonucleotides 3243_Trg-tetR and 3244_trg-tetA

B: TH14977 (Δ*flgMN7753* Δ*flgKL7770* Δ*trg-7722*::*tetRA*)

C: TH14735 (Δ*trg-7774*); made using oligonucleotides 3410_Trgdelfwd and3411_Trgdelrev

D: TH14978 (Δ*flgMN7753* Δ*flgKL7770* Δ*trg-7774*)

Starting strain: TH14978

A: TH14589 (Δ*ycgR7751*::*tetRA*); made using oligonucleotides 3255_YcgRtetRand 3256_YcgRtetA

B: TH14979 (Δ*flgMN7753* Δ*flgKL7770* Δ*trg-7774* ΔycgR7751::*tetRA*)

C: TH14779 (Δ*ycgR7775*); made using oligonucleotides 3424_YcgRcleandelfwd and 3425_YcgRDELREV

D: TH15060 (Δ*flgMN7753* Δ*flgKL7770* Δtrg-7774 Δ*ycgR7775*)

Starting strain: TH15060

A: TH14238 (Δ*fliB-T7727*::*tetRA*); made using oligonucleotides 3237_FliB-FliT-tetR and 3238_fliB-FliT-tetA

B: TH15061 (Δ*flgMN7753* Δ*flgKL7770* Δ*trg-7774* Δ*ycgR7775* Δ*fliB-T7727*::*tetRA*)

C: TH14691 (Δ*fliB-T7771*); made using oligonucleotides 3404_FliB-FliTdelfwd and 3405_FliB-FliTdelrev

D: TH15138 (Δ*flgMN7753* Δ*flgKL7770* Δ*trg-7774* Δ*ycgR7775* Δ*fliB-T7771*)

Starting strain: TH15138

A: TH14301 (Δ*hin-fljA7731*::*tetRA*); made using oligonucleotides 3231_FljA-hin-tetR and 3232_FljA-hin-tetA

B: TH15201 (Δ*flgMN7753* Δ*flgKL7770* Δ*trg-7774* Δ*ycgR7775* Δ*fliB-T7771* Δ*hin-fljA7731*::*tetRA*)

C: TH14591 (Δ*hin-fljA7752*); made using oligonucleotides 3427_FljAdelrwd and 3428_FljAdelrev

D: TH15202 (Δ*flgMN7753* Δ*flgKL7770* Δ*trg-7774* Δ*ycgR7775* Δ*fliB-T7771* Δ*hin-fljA7752*)

Starting strain: TH15202

A: TH14237 (Δ*prgH-hilDA7726*::*tetRA*); made using oligonucleotides 3253_PrgH-HilA-tetR and 3254-PrgH-HilA-tetA

B: TH15203 (Δ*flgMN7753* Δ*flgKL7770* Δ*trg-7774* Δ*ycgR7775* Δ*fliB-T7771* Δ*hin-fljA7752* Δpr*gH-hilA7726*::*tetRA*)

C: TH14864 (Δ*prgH-hilDA7791*); made using oligonucleotides 3422_PrgH-HilA-DELFWD and 3423_PrgH-HilA-DEL REV

D: TH15204 (Δ*flgMN7753* Δ*flgKL7770* Δ*trg-7774* Δ*ycgR7775* Δ*fliB-T7771* Δ*hin-fljA7752* Δ*prgH-hilA7791*)

Starting strain: TH15204

A: TH14236 (Δ*cheV7725*::*tetRA*); made using oligonucleotides 3229_cheV-tetR and 3230_chev-tetA

B: TH15313 (Δ*flgMN7753* Δ*flgKL7770* Δ*trg-7774* Δ*ycgR7775* Δ*fliB-T7771* Δ*hin-fljA7752* Δ*prgH*-*hilA7791* Δ*cheV7725*::*tetRA*)

C: TH15173 (Δ*cheV7829*); made using oligonucleotides 3418_STM2314delfwd and 3419_STM2314delrev

D: TH15314 (Δ*flgMN7753* Δ*flgKL7770* Δ*trg-7774* Δ*ycgR7775* Δ*fliB-T7771* Δ*hin-fljA7752* Δ*prgH-hilA7791* Δ*cheV7829*)

Starting strain: TH15314

A: TH14305 (Δ*mcpA*(*STM3138*)*-7733*::*tetRA*); made using oligonucleotides 2410_McpA-tetR and 2411_McpA-tetA

B: TH15315 (Δ*flgMN7753* Δ*flgKL7770* Δ*trg-7774* Δ*ycgR7775* Δ*fliB-T7771* Δ*hin-fljA7752* Δ*prgH-hilA7791* Δ*cheV7829* Δ*mcpA7733*::*tetRA*)

C: TH14868 (Δ*mcpA7792*); made using oligonucleotides 3416_STM3138delfwd and 3417_STM3138delrev

D: TH15316 (Δ*flgMN7753* Δ*flgKL7770* Δ*trg-7774* Δ*ycgR7775* Δ*fliB-T7771* Δ*hin-fljA7752* Δ*prgH-hilA7791* Δ*cheV7829* Δ*mcpA7792*)

Starting strain: TH15316

A: TH14228 (Δ*tcp-7721*::*tetRA*); made using oligonucleotides 3241_Tcp-tetR and 3242_Tcp-tetA

B: TH15398 (Δ*flgMN7753* Δ*flgKL7770* Δ*trg-7774* Δ*ycgR7775* Δ*fliB-T7771* Δ*hin-fljA7752* Δ*prgH-hilA7791* Δ*cheV7829* Δ*mcpA7792* Δ*tcp-7721*::*tetRA*)

C: TH15171 (Δ*tcp-7827*); made using oligonucleotides 3408_tcpdelfwd and 3409_tcpdelrev

D: TH15399 (Δ*flgMN7753* Δ*flgKL7770* Δ*trg-7774* Δ*ycgR7775* Δ*fliB-T7771* Δ*hin-fljA7752* Δ*prgH-hilA7791* Δ*cheV7829* Δ*mcpA7792* Δ*tcp-7827*)

Starting strain: TH15399

A: TH14239 (Δ*yhjH7728*::*tetRA*); made using oligonucleotides 3399_YhjHtetA and 3400_YhjHtetR

B: TH15400 (Δ*flgMN7753* Δ*flgKL7770* Δ*trg-7774* Δ*ycgR7775* Δ*fliB-T7771* Δ*hin-fljA7752* Δ*prgH-hilA7791* Δ*cheV7829* Δ*mcpA7792* Δ*tcp-7828* Δ*yhjH7728*::*tetRA*)

C: TH14554 (Δ*yhjH7740*); made using oligonucleotides 3420_YhjHDELRWD and 3421_YhjHDELREV

D: TH15401 (Δ*flgMN7753* Δ*flgKL7770* Δ*trg-7774* Δ*ycgR7775* Δ*fliB-T7771* Δ*hin-fljA7752* Δ*prgH-hilA7791* Δ*cheV7829* Δ*mcpA7792* Δ*tcp-7828* Δ*yhjH7740*)

Starting strain: TH15401

A: TH14226 (Δ*tsr-7720*::*tetRA*); made using oligonucleotides 3239_Tsr-tetR and 3240_Tsr-tetA

B: TH15653 (Δ*flgMN7753* Δ*flgKL7770* Δ*trg-7774* Δ*ycgR7775* Δ*fliB-T7771* Δ*hin-fljA7752* Δ*prgH-hilA7791* Δ*cheV7829* Δ*mcpA7792* Δ*tcp-7827* Δ*yhjH7740* Δ*tsr-7720*::*tetRA*)

C: TH15172 (Δ*tsr-7828*); made using oligonucleotides 3406_Tsrdelfwd and 3407_tsrdelrev

D: TH15654 (Δ*flgMN7753* Δ*flgKL7770* Δ*trg-7774* Δ*ycgR7775* Δ*fliB-T7771* Δ*hin-fljA7752* Δ*prgH-hilA7791* Δ*cheV7829* Δ*mcpA7792* Δ*tcp-7827* Δ*yhjH7740* Δ*tsr-7828*)

Starting strain: TH15654

A: TH14232 (Δ*mcpB*(*STM3152*)-7723::*tetRA*); made using oligonucleotides 3245_STM3152-tetR and 3246_STM3152-tetA

B: TH15655 (Δ*flgMN7753* Δ*flgKL7770* Δ*trg-7774* Δ*ycgR7775* Δ*fliB-T7771* Δ*hin-fljA7752* Δ*prgH-hilA7791* Δ*cheV7829* Δ*mcpA7792* Δ*tcp-7827* Δ*yhjH7740* Δ*tsr-7828* Δ*mcpB7723*::*tetRA*)

C: TH15357 (Δ*mcpB7835*(*STM3152*)); made using oligonucleotides 3412_STM3152delfwd and 3413_STM3152delrev

D: TH15656 (Δ*flgMN7753* Δ*flgKL7770* Δ*trg-7774* Δ*ycgR7775* Δ*fliB-T7771* Δ*hin-fljA7752* Δ*prgH-hilA7791* Δ*cheV7829* Δ*mcpA7792* Δ*tcp-7827* *ΔyhjH7740* Δ*tsr-7828* Δ*mcpB7835*)

Starting strain: TH15656

A: TH14234 (Δ*aer-mcpC*(*STM3216*)-*7724*::*tetRA*); made using oligonucleotides 3247_Aer-tetR and 3248_Aer-tetA

B: TH15697 (Δ*flgMN7753* Δ*flgKL7770* Δ*trg-7774* Δ*ycgR7775* Δ*fliB-T7771* Δ*hin-fljA7752* Δ*prgH-hilA7791* Δ*cheV7829* Δ*mcpA7792* Δ*tcp-7827* Δ*yhjH7740* Δ*tsr-7828* Δ*mcpB7835* Δ*aer-mcpC7724*::*tetRA*)

C: TH15356 (Δ*aer-mcpC7834*(*STM3216*)); made using oligonucleotides 3414_Aerdelfwd and 3415_aerdelrev

D: TH15698 (Δ*flgMN7753* Δ*flgKL7770* Δ*trg-7774* Δ*ycgR7775* Δ*fliB-T7771* Δ*hin-fljA7752* Δ*prgH-hilA7791* Δ*cheV7829* Δ*mcpA7792* Δ*tcp-7827* Δ*yhjH7740* Δ*tsr-7828* Δ*mcpB7835* Δ*aer-mcpC7834*)

Starting strain: TH15698

A: TH15143 (P*_motAB_7795* Δ*motA-cheZ7809*::*tetRA*); made using oligonucleotides 3261_MotA-CheZ-tetR and 3262_MotA- CheZ-tetA

B: TH15902 (Δ*flgMN7753* Δ*flgKL7770* Δ*trg-7774* Δ*ycgR7775* Δ*fliB-T7771* Δ*hin-fljA7752* Δ*prgH-hilA7791* Δ*cheV7829* Δ*mcpA7792* Δ*tcp-7827* Δ*yhjH7740* Δ*tsr-7828* Δ*mcpB7835* Δ*aer-mcpC7834* P*_motAB_7795* Δ*motA-cheZ7809*::*tetRA*)

C: TH15901 (P*_motAB_7795* Δ*motA-cheZ7888*); made using oligonucleotides 3263_MotA-CheZclean and 3264_PmotABfillin

D: TH15903 (Δ*flgMN7753* Δ*flgKL7770* Δ*trg-7774* Δ*ycgR7775* Δ*fliB-T7771* Δ*hin-fljA7752* Δ*prgH-hilA7791* Δ*cheV7829* Δ*mcpA7792* Δ*tcp-7827* Δ*yhjH7740* Δ*tsr-7828* Δ*mcpB7835* Δ*aer-mcpC7834* P*_motAB_7795* Δ*motA-cheZ7888*)

Lastly, strain TH22673 (Δ*fliA5805*::*tetRA* Δ*flgMN7753* Δ*flgKL7770* Δ*trg-7774* Δ*ycgR7775* Δ*mcpA7792* Δ*tsr-7828* Δ*mcpB7835* Δ*hin-fljA7752* Δ*prgH-hilA7791* Δ*cheV7829* Δ*tcp-7827* Δ*yhjH7740* Δ*aer-mcpC7834* P*_motA_7795* Δ*motA-cheZ7888*) was constructed by P22 transduction and Tc^R^ selection of Δ*fliA5805*::*tetRA* *fliB5470*::Mu*d*K from TH7410 (Δ*fliA5805*::*tetRA* *fliB5470*::Mu*d*K; Hughes lab strain collection) into TH15903.
